# EEG Analyses of visual cue effects on executed movements

**DOI:** 10.1101/2024.04.22.590535

**Authors:** Patrick Suwandjieff, Gernot R. Müller-Putz

**Affiliations:** Institute of Neural Engineering, Graz University of Technology, Graz, Austria; BioTechMed Graz, Austria

**Keywords:** brain-computer interface (BCI), visual cue, visual evoked potentials, electroencephalogram (EEG), movement-related cortical potential (MRCP), cue-based, self-paced

## Abstract

**Background:** In electroencephalographic (EEG) or electrocorticographic (ECoG) experiments, visual cues are commonly used for timing synchronization but may inadvertently induce neural activity and cognitive processing, posing challenges when decoding self-initiated tasks.

**New Method:** To address this concern, we introduced four new visual cues (Fade, Rotation, Reference, and Star) and investigated their impact on brain signals. Our objective was to identify a cue that minimizes its influence on brain activity, facilitating cue-effect free classifier training for asynchronous applications, particularly aiding individuals with severe paralysis.

**Results:** 22 able-bodied, right-handed participants aged 18-30 performed hand movements upon presentation of the visual cues. Analysis of time-variability between movement onset and cue-aligned data, grand average MRCPs, and classification outcomes revealed significant differences among cues. Rotation and Reference cue exhibited favorable results in minimizing temporal variability, maintaining MRCP patterns, and achieving comparable accuracy to self-paced signals in classification.

**Comparison with Existing Methods:** Our study contrasts with traditional cue-based paradigms by introducing novel visual cues designed to mitigate unintended neural activity. We demonstrate the effectiveness of Rotation and Reference cue in eliciting consistent and accurate MRCPs during motor tasks, surpassing previous methods in achieving precise timing and high discriminability for classifier training.

**Conclusions:** Precision in cue timing is crucial for training classifiers, where both Rotation and Reference cue demonstrate minimal variability and high discriminability, highlighting their potential for accurate classifications in online scenarios. These findings offer promising avenues for refining brain-computer interface systems, particularly for individuals with motor impairments, by enabling more reliable and intuitive control mechanisms.

## Introduction

Visual cues are commonly used during electroencephalographic (EEG) or electrocorticographic (ECoG) experiments to ensure that the timing of the recorded brain signals is synchronized to specific events (e.g., the execution of a specific hand movement). However, these cues can elicit neural activity (e.g., visual evoked potentials (VEPs), cognitive processing) that can obscure the neural dynamics associated with other brain signals, such as those arising from movement (Schwarz et al. 2018; Scheel et al. 2015; Ofner et al. 2019; Pearce et al. 2017). VEPs are electrical responses generated by the brain in response to visual stimuli. When a person is exposed to a visual stimulus, such as a flashing light or a patterned image, the brain’s neural activity synchronizes and produces electrical potentials that can be detected from the surface of the brain and the scalp (via EEG). VEPs provide valuable insights into visual processing, offering a non-invasive means to study the timing and characteristics of neural responses related to visual stimuli (Odom et al. 2004). While informative for examining visual processing, they can pose challenges when analyzing other brain signals, for instance movement-related cortical potentials (MRCPs). MRCPs, as described by (Kornhuber and Deecke 1965; Shibasaki and Hallett 2006), are brain signals linked with the preparation and execution of voluntary movements as well during imagination and attemption. These signals encompass components such as the Bereitschaftspotential (BP), which precedes movement onset, the motor potential (MP), that occurs during the actual execution of the movement and the movement-monitoring potential (MMP) which occurs thereafter. These events are thought to correspond to movement planning and preparation, execution, and the control of performance, respectively and are usually revealed by locking the low-frequency EEG to the movement onset. MRCPs are known for encoding various elements of movement, such as movement directions (Kobler, Sburlea, et al. 2020; Kobler, Kolesnichenko, et al. 2020; Pereira et al. 2017; Waldert et al. 2009), grasp types (Schwarz et al. 2018; Jochumsen et al. 2016), speed (Gu, Dremstrup, and Farina 2009) or force (Gu, Dremstrup, and Farina 2009; Jochumsen et al. 2013). The challenge of cues eliciting neural activity becomes particularly pronounced when decoders, trained on cue-based paradigms, are applied to identify or classify self-initiated movements from low-frequency EEG. This ‘cue effect’ was investigated by (G. Pfurtscheller et al. 2008) in which the central focus layed on the brain’s response to a visual cue. The authors utilized various neuroimaging techniques such as EEG and functional magnetic resonance imaging (fMRI) to measure cortical activity during motor imagery following a cue. The results of the study revealed a short-lived yet significant change in brain activity immediately following the cue. Such changes indicate that the brain responds to the cue and prepares or initiates the mental imagery of the action. This finding suggests that cues play a crucial role in priming the brain for subsequent motor imagery tasks, potentially facilitating the generation of mental representations of actions. Efforts to mitigate the impact of the visual cue on MRCPs were made by (Ofner et al. 2019) through the implementation of a novel cue design. In this design, the visual presentation of the cue gradually diminished over time instead of abruptly indicating the start of the movement. By adjusting the stimulus in such a way, the occurrence of VEPs was diminished, resulting in less pronounced influence on MRCPs. It can be said that the use of a visual cue is critically important for effective brain-computer interface (BCI) operation in locked-in syndrome (LIS) patients since there is no way to detect the actual onset of an attempted movement. Furthermore, previous studies have demonstrated the persistence of the cue effect at the ECoG level (Branco et al. 2017). As previously noted this interference becomes a concern when applying decoders trained on cue-based paradigms to classify or identify movements or movement attempts, as the neural dynamics related to the visual cues can obscure the nuanced patterns of signals like MRCPs. Careful consideration is required to disentangle the distinct contributions of visual processing and movement-related activity in such experimental settings. Meaning, within the realm of BCIs, asynchronous configurations present challenges to decoding performance due to the classifier being trained on MRCPs influenced by cue-related potentials, which are not present in asynchronous usage. Consequently, it is essential to train a classifier using MRCPs minimally influenced by the cue. Therefore, the primary objective of this study was to address a fundamental question: is it possible to create a visual cue that has no or minimal influence on MRCPs in order to find a cue beneficial for the usage of decoding movement attempts? While existing research has already compared cue-based and self-paced MRCPs (Savić et al. 2014) respectively visual, tactile and auditory cues (Cincotti et al. 2007; Scheel et al. 2015; Pearce et al. 2017), there has been limited exploration into developing strategies to mitigate the influence of visual cues on MRCPs. To address this void, we draw inspiration from prior research conducted by (Ofner et al. 2019) and (Gert Pfurtscheller et al. 2010). Building on this foundation, we devise three new visual cues with the underlying principle that their properties would exert minimal influence on signals connected with movement execution. By rejecting or minimizing cue-induced effects, our aim is to ensure that signals recorded during cue-based activities closely mirror those generated during self-initiated movement attempts. This optimization seeks to elevate decoding performance and facilitate a more seamless control of BCI. Moreover, it enables cue-effect free classifier training for subsequent asynchronous applications. This study could aid in achieving stable and robust four-direction discrete cursor control, thereby enhancing user freedom. To fulfill these objectives, we employed four distinct gestures which were chosen based on classification results of previously performed studies (Ofner et al. 2019; Pereira, Sburlea, and Müller-Putz 2018; Ofner et al. 2017).

## Materials and methods

### Participants

In this study, a total of 22 healthy participants (12 male, 10 female) with an average age of 26.2 ± 4.2, engaged in the recording of EEG activity. All participants provided written informed consent after receiving comprehensive information from the experimenters regarding the study’s objectives, content, and procedures. Participants retained the right to terminate or discontinue their participation at any time without providing a reason and without facing any consequences. The experimental study received approval from the local ethics committee at TU Graz before commencement. Furthermore, the data recorded for each participant underwent anonymization.

### Experimental Design

The experiment consisted of two parts: (i) cue-based data collection applying four different visual cues and four different gestures. (ii) self-paced phase where participants were instructed to freely execute the corresponding four gestures at any time.

#### Visual Cues

The cues were designed with the intention of minimizing any potential impact on brain signals resulting from visual stimuli. As a result, it is crucial to avoid rapid changes during the presentation. The core concept underlying these cues is their gradual appearance; they do not appear abruptly but transition smoothly to their initial positions through methods like shrinking, rotating, or fading.

The Reference Cue, selected based on the findings of the study by (Ofner et al. 2019), comprises a green filled circle that initiates a random-speed shrinking process. Upon the shrinking circle reaching the circumference of a small white circle (representing the go cue), participants are prompted to execute the movement.

The Fading Cue comprises a white fixation cross surrounded by a green circle. Over time, it gradually transitions to match the color of the background, and the execution of the movement is intended to coincide with the cue fully matching the background color.

In the Rotation Cue, two white fixation crosses are displayed, with one of them rotated by ±45 degrees. The rotated fixation cross initiates a continuous turning motion, and participants are prompted to perform a movement when its position aligns with the non-rotated cross. The uniqueness of this cue lies in the sustained Rotation movement throughout the entire stimulus presentation.

The Star Cue, a white fixation cross is presented against a black background, and its edges form a Star shape. The Star gradually diminishes as the four central point’s move closer to the origin of the fixation cross. Participants are instructed to execute the movement when these points completely overlap.

The initial, approximately middle, and final positions of each cue can be observed in Fig. 1.

**Figure 1:**
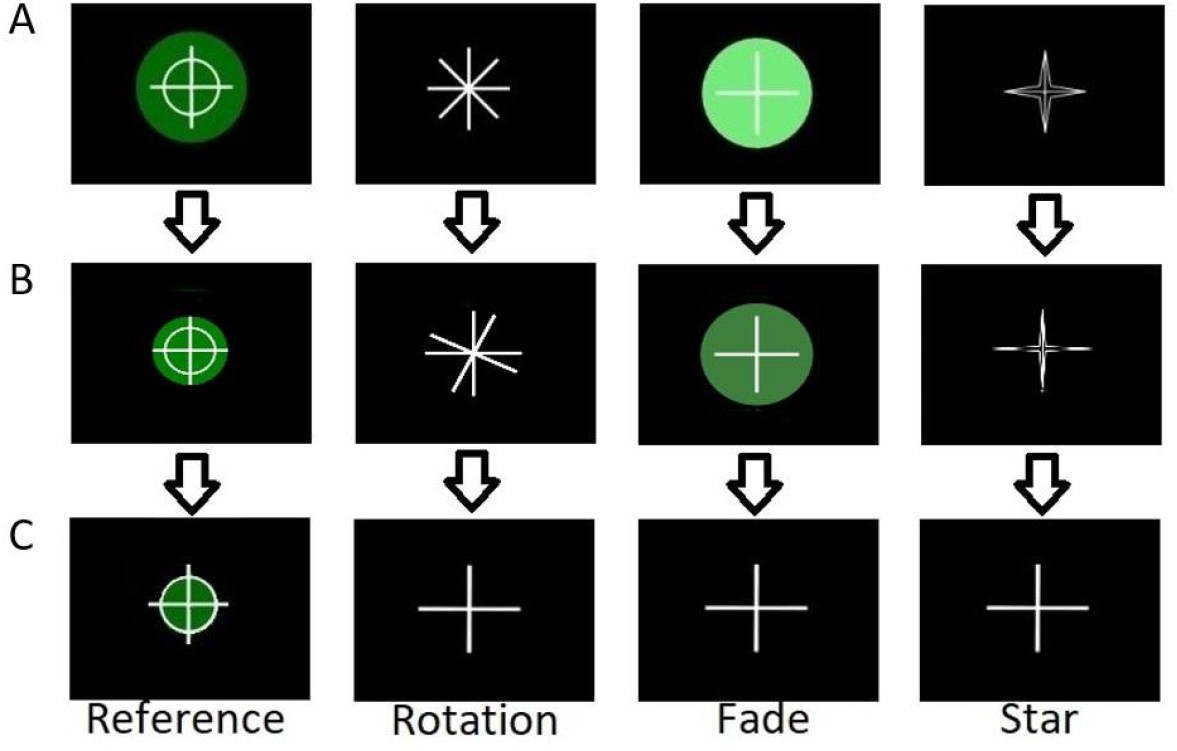
Cues working principle. A) displays the initial position of the cue type Reference, Rotation, Fading, Star. B) shows the middle position of the cues. C) illustrates the final position.

#### Paradigm

For the cue-based part participants were instructed to execute four specific gestures and rest precisely at predetermined start times, indicated by the four distinct visual cues introduced in the preceding section. The initial position before the execution of a specific gesture was termed ‘rest’ and participants consistently returned to this neutral starting position after performing a gesture. This initial position was used as a 5th ‘gesture’. Additionally the following gestures were executed: pincer grasp, pistol, fist, and ‘Y’-gesture of the American sign language. These four gestures and the rest condition had to be performed six times (randomly ordered) within one run. For the rest condition participants just stayed in the neutral starting position. The duration of such a run was 5 min, it was followed by a 30 s rest period. The type of cue was constant throughout one run. This process was repeated over 32 separate runs, each with a randomly selected cue, resulting in a cumulative total of 48 trials per gesture and per cue. A single trial, exemplified on the Rotation Cue, has the following sequence (Fig. 2): the gesture was displayed for 1 s, followed by two crosses with one of them rotated by ± 45 degrees (‘ready phase’) which remained in their position for 0.5 - 1 s. The rotated cross started to rotate and it took 2 - 3 s that their positions were identical (‘preparation phase’). This phase was designed to function as a preparatory period for participants, serving as a smooth visual transition between cues and minimizing visual cue effects on EEG. After this, the participants had to execute the movement (3 s of execution and holding the end position of the gesture). This phase is called the ‘go’ phase. After that a ‘rest’ phase with a blank screen was presented for 1.5 s. The other cues followed a similar timing.

**Figure 2:**
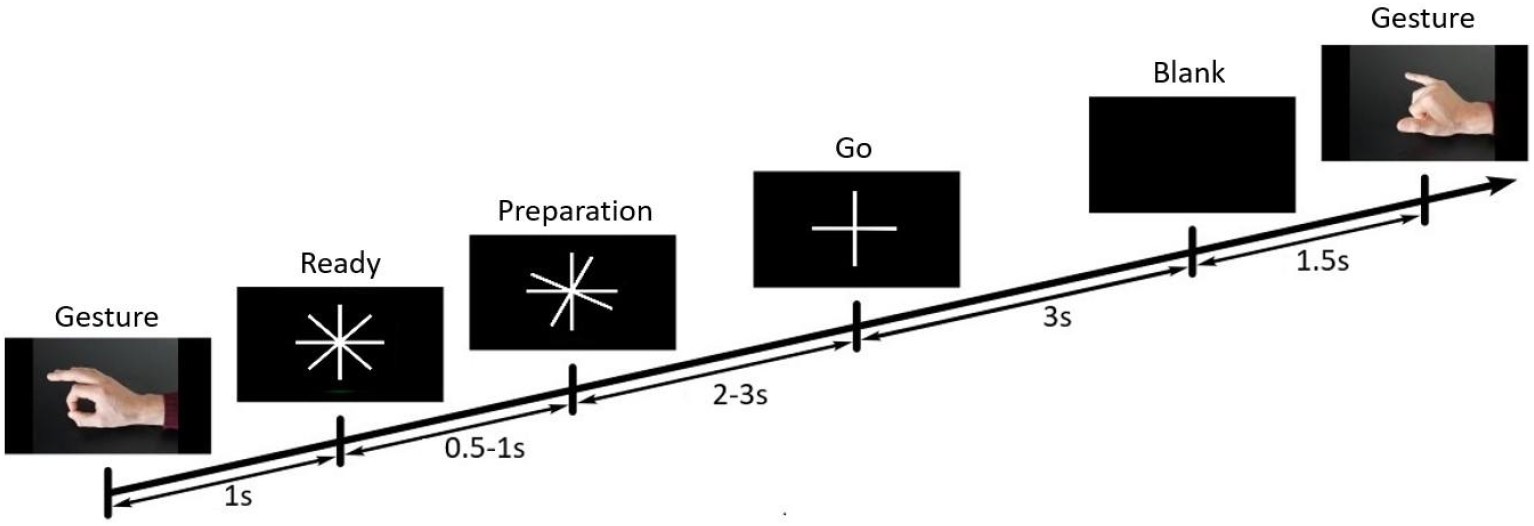
Timings of the different phases during one gesture trial. Starting with the gesture presentation, followed by the ready phase of the cue, thereafter the preparation phase, then the go phase and at last the blank (rest) phase.

In the self-paced part, participants were instructed to execute the same gesture (which was shown for 1.5 s at the start) and hold it for 3 s (same procedure as in the cue-based part) at approximately 10 s intervals over the course of a 5-min run. The timing (including the instruction of which gesture to use) of one run can be seen in Fig. 3. This protocol was repeated across a total of 8 runs, leading to 60 trials for each distinct gesture. It is worth mentioning that the amount of movement trials for each subject differ slightly, since the self-paced paradigm instructs the participants to do the movements approximately every 10 s, leading to some variation between subjects.

**Figure 3:**
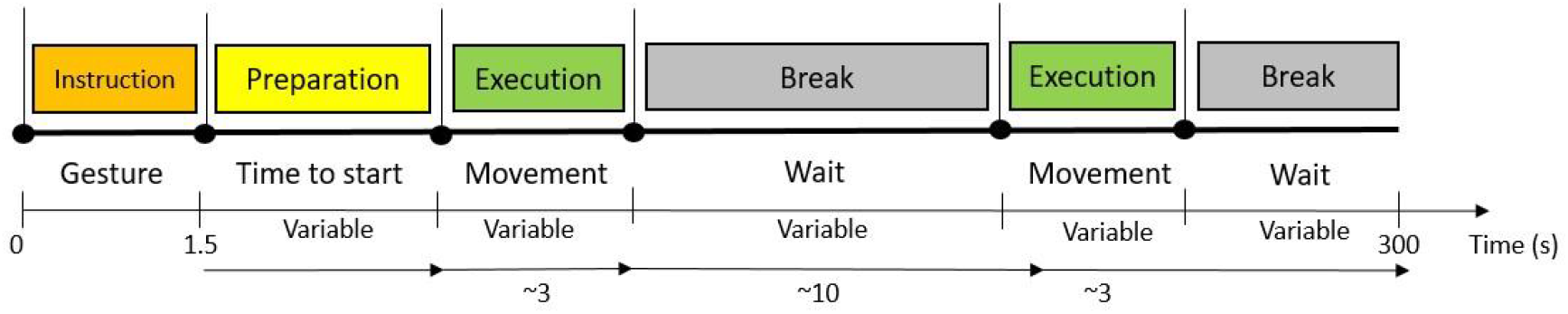
Paradigm & experimental timeline of a 5-min run in the self-paced segment.

### Signal Recording

The EEG signals were captured using a 64-channel actiCAP system (Brain Products GmbH, Gilching, Germany) with a sampling rate of 500 Hz. The electrodes were arranged based on the 10-10 electrode system, encompassing 60 electrodes covering all cortical areas. Additionally, four electrodes were dedicated to electrooculography (EOG) purposes. These EOG electrodes were strategically positioned at the outer canthi of both eyes and above and below the left eye to monitor saccades and blinks. The ground electrode was located at the right mastoid, while the Reference electrode was placed at FCz.

To detect actual movement onsets in both cue-based and self-paced sessions, we employed a custom-built motion capture system with a sampling frequency of 30 Hz. A marker was placed at the tip of the participant’s index finger. The data provided spatial information along the x, y, and z axes.

Both data streams were synchronized with the help of the lab streaming layer (LSL).

#### Signal Preprocessing

The signals recorded underwent processing and analysis utilizing MATLAB R2019a alongside the EEGLAB toolbox ([16]). The 60-channel EEG signals were initially zero-phase band-pass filtered (Butterworth filter of 3rd order) between 0.5 and 70 Hz. 50 Hz power line interference was removed using a notch filter. Eye and muscle artifacts were removed through independent component analysis (ICA). Thereafter, common average reference (CAR) was applied. As our focus in this study is specifically on MRCPs, we applied bandpass filtering to the data within the low-frequency range of (0.5 - 5 Hz) using a 3rd order butterworth IIR filter. Subsequently, we resampled the entire signal at 20 Hz to reduce computational workload. The resampling included the application of an antialiasing filter to prevent edge effects. All trials were temporally aligned and epoched either with respect to cue onset of the movement onset, within a window spanning from -2 s to 2 s.

The kinematic data of gestures was employed to calculate the velocity of the participant’s hand movements. Movement onset was identified when the hand’s velocity exceeded a predefined threshold (which was set to the same velocity value for all participants) during the interval between the ‘go phase’ and the cue for the break, ensuring precise detection while avoiding false positives for small movements during rest.

The dataset of one participant had to be excluded because of bad signal quality and kinematic tracking.

#### Timing Variability between cue alignment and movement onset

We investigated the temporal variability between the cue onset and the actual initiation of movement by the participants, referred to as the movement onset. This analysis considered all trials encompassing every movement by each participant for every cue. The timing variability was defined as follows: For each executed movement, the exact initiation time, determined through motion capture, was subtracted from the time indicated by the cue. The objective was to assess the performance of each cue in terms of its temporal variability, with a precise indication of movement onset being the desired outcome.

#### Analysis of MRCPs

We analyzed the EEG signals in the frequency range of 0.5 - 5 Hz to examine MRCPs. The grand average for all participants and gestures was used to calculate MRCPs in the time domain for each cue. Our focus in the time-domain analysis centered on electrode C1, given the right-hand execution, expecting the MRCPs to manifest contralaterally. In the cue-based segment of the study, our analysis concentrated on both cue-aligned data and movement onset. This approach allowed us to discern differences in MRCPs influenced by the cue and those unaffected by temporal variability respectively VEPs induced from the cue. Our primary focus was on comparing the grand average MRCPs generated by the four cues with the grand average MRCPs from the self-paced segment of the study. Therefore we performed as well a Wilcoxon ranksum test and corrected for multiple comparisons using the Benjamini-Hochberg method to examine potential variations in results among the different cue triggered MRCPs compared to the self-paced MRCPs. Furthermore, we generated topographical maps for each cue, triggering the analysis on both cue onset and movement onset. These maps were then compared with the self-paced data, where the analysis was triggered on movement onset. This comparison aimed to highlight distinct processing patterns and MRCP distributions across the scalp.

#### Classification - movement vs rest

For the classification, we combined all gestures (‘movement’) and contrasted them against the rest state (‘no movement’) for each cue (cue aligned & movement onset aligned) respectively as well for self-paced (movement onset aligned). We therefore merged, for every participant, trials from all four gestures into a single class, labeled as ‘movement’ (4 gestures x 48 trials = 192 trials for each cue type; 4 gestures x 60 = 240 trials for self-paced). Trials from the rest condition (independently of the cue type) were utilized as a ‘rest’ class (4 cues x 48 trials = 192 trials). To identify the point of maximum discrimination between the two classes around the cue onset, we employed a 2-class shrinkage linear discriminant analysis (sLDA) (Peck and Van Ness 1982; Blankertz et al. 2011), using one sample overlapping 1-s window segments which were shifted along the epochs within each participant (compare (Ofner et al. 2019)). The classification model was constructed based on a 10×1 fold cross-validation scheme.

## Results

### Time-variability between cue alignment and movement onset

In Fig. 4 (A), the distinctively narrower distribution of the Rotation and Reference cues around time 0 s is clearly evident. The timing variability has been normalized to correct for a constant offset. Notably, the number of trials for the Rotation and Reference cues slightly exceeds 500, while for the Fade and Star cues, it remains at 400 trials around time 0 s. Additionally It is clearly visible in Fig.4 (B) that the shape of the timing variability of the Rotation cue and the Reference cue distinctly indicates that the values for these two cues are more centered around the 0 s point (matching point for movement onset and cue alignment), with a slight offset in the negative time direction. Conversely, the Fade cue and the Star cue exhibit a much broader distribution. The average time differences between movement onset and cue onset for different cues are as follows: registering the shortest delay at -0.11 s is the Rotation cue, followed by the Reference cue with a delay of -0.13 s, then the Fade cue at -0.15 s, and finally, the Star cue with the lengthiest delay at -0.19 s. On average, participants tend to execute the gesture a bit earlier than the cues indicated to them to do so. The statistical difference test reveals no significant distinction between the outcomes of the Rotation and Reference cue. However, for all other scenarios, a significant difference is observed.

**Figure 4:**
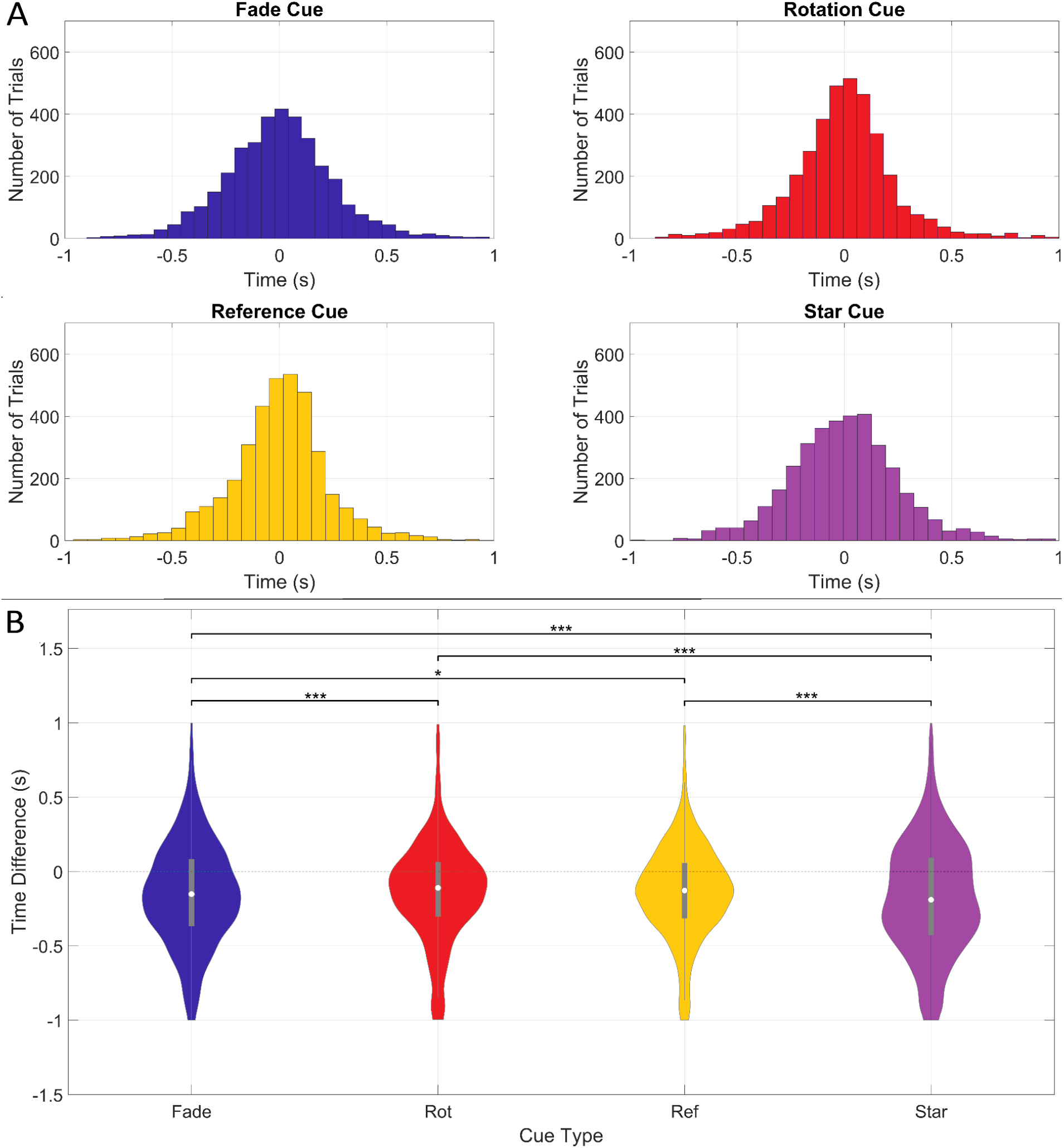
A) Histogram of the normalized time difference between movement onset and cue aligned for the different cue types. B) Violin plot of the time difference between movement onset and cue aligned for the different cue types. Statistically significant differences between cues are indicated with stars (*p<0.05, **p<0.01, ***p<0.001). A and B: Fade Cue (blue), Rotation Cue (red), Reference Cue (yellow), Star Cue (magenta).

### Grand average MRCPs time signal

The grand average differences among MRCPs (C1) aligned to the cue onset, as depicted in Fig. 5 (B), distinctly highlight that the configurations of the Rotation and Reference cue produce a pronounced MRCP shape while MRCPs following the Fade and Star cue show weak and undefined shapes. Specifically, the Rotation cue exhibits a slightly higher positive peak during the movement-monitoring potential compared to the Reference cue, while the timing of its negative peak aligns more closely to the time 0 s (cue aligned). The Fade and Star cue produce a much more varying result. It is notable that the BP occurs the earliest in the MRCPs generated by the Star cue. When comparing the MRCPs from the different cues aligned to the movement onset (Fig. 5 (A)), it is clearly visible that they have nearly the same form. The shape of the MRCPs aligned to the movement onset is much more pronounced to those aligned to the cue onset. The Fade cue seems to have the most influence, while the other cues have no visible difference during the time between -2 to -1.5 s of the preparation phase. Additionally, we conducted a comparison between the grand average MRCPs of the four cues and the grand average MRCPs of the self-paced recordings, as illustrated in Fig. 6. Reference and Rotation cue closely resemble the MRCP shape observed in the self-paced approach. In contrast, the MRCP shapes corresponding to the Fade and Star cue have nearly no similarity at all. The statistical significance of the similarity at each time point between the MRCPs of the cues and those of the self-paced data is illustrated to underscore the observed variations. This analysis provides a comprehensive understanding of the distinct features present in the MRCPs elicited by different cues compared to the self-paced data. It is noteworthy that the Rotation, Reference, and Star Cue exhibit similar levels of similarity to the self-paced approach until the onset of the motor potential (MP), with their levels being visible higher compared to the results of the Fade Cue. Conversely, the Fade Cue demonstrates the highest similarity after the MP among all cue types.

**Figure 5.**
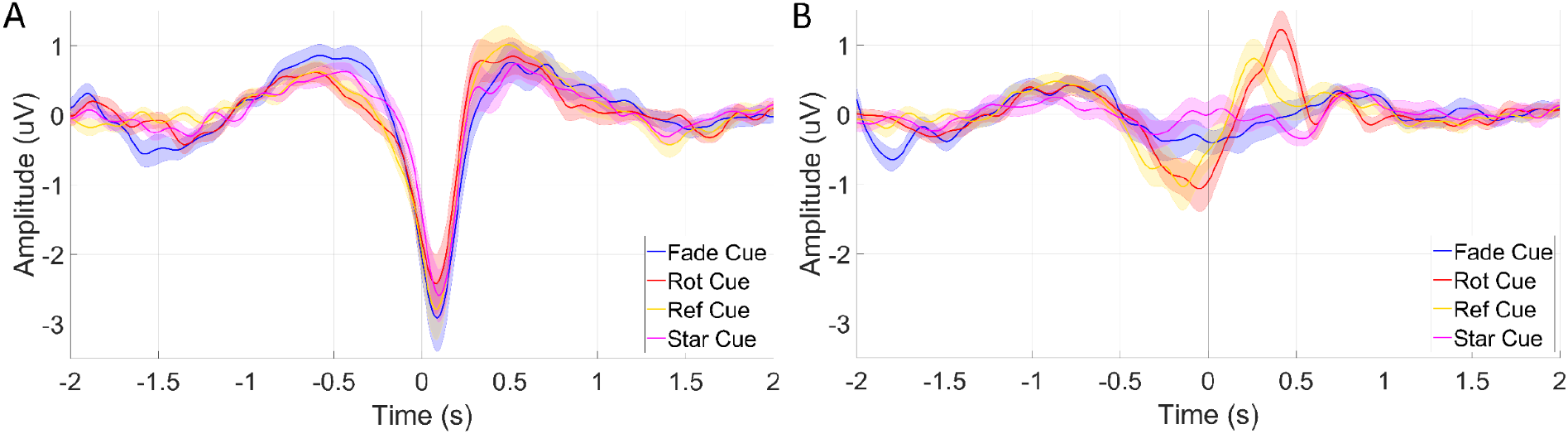
A) Cue-Specific MRCPs (C1) cue aligned (mean ± standard error). B) Cue-Specific MRCPs (C1) movement aligned (mean ± standard error). Each MRCP is assigned a specific color: Fade Cue (blue), Rotation Cue (red), Reference Cue (yellow), Star Cue (magenta). The timescale is aligned to onset for both panels.

**Figure 6.**
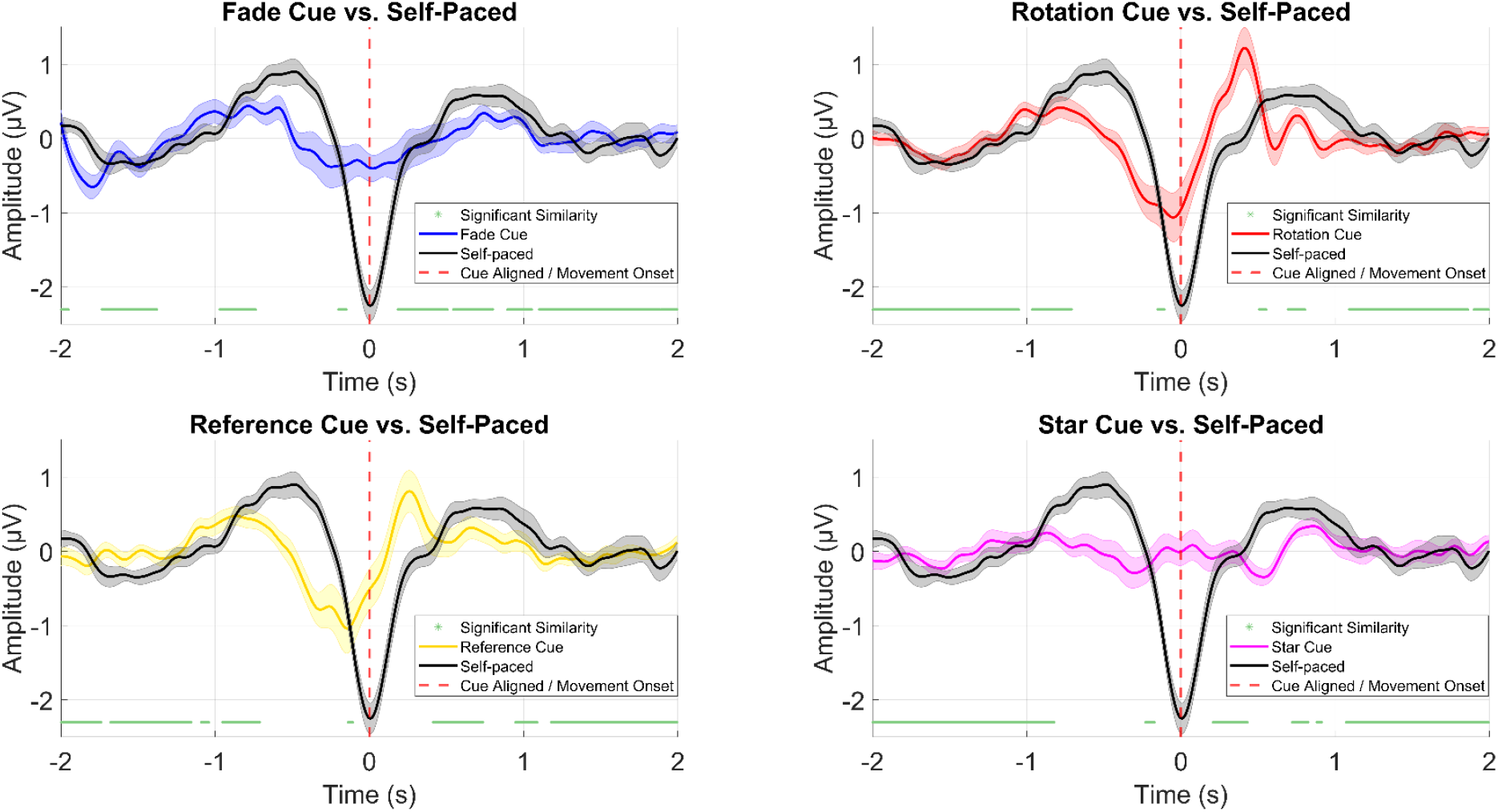
Time-Domain Differences Between Individual Cues and Self-Paced Data of Electrode C1 (± standard error). Fade Cue (blue), Rotation Cue (red), Reference Cue (yellow), Star Cue (magenta), self-paced (black) and significant differences (green) at every time point (p<0.05). The red dashed line at t = 0 denotes the cue onset respectively movement onset.

### Grand average MRCPs Topoplots

The spatial map results reveal a significant distinction between the self-paced data, triggered by movement onset, and the cue-based data, aligned to cue onset. Notably, in Fig. 7, the Star cue data displays the least pronounced positive amplitude before movement execution (−1 to 0 s), with a more prominent negative slope observed on the right hemisphere of the scalp (0s). Conversely, the remaining three cue-aligned patterns demonstrate comparable behavior within the -1 s to 0 s timeframe. Specifically, the Reference and Rotation Cue patterns exhibit dominant positive amplitudes at timepoints 0.25 s and 0.375 s, respectively. These patterns are more pronounced in the occipital area for the Reference cue and in the central region for the Rotation Cue. In contrast, the findings from movement onset triggered cue-based data reveal a similar pattern across all cue produced MRCPs, as depicted in Fig. 8. Notably, the MRCPs influenced visually by the Star cue exhibit a distinctly different pattern generated at -1 s and 1 s compared to the other three cue generated data. The self-paced spatial map indicates generally lower activity for all time points, with its negative slope predominantly generated in the motor area and on the left hemisphere.

**Figure 7:**
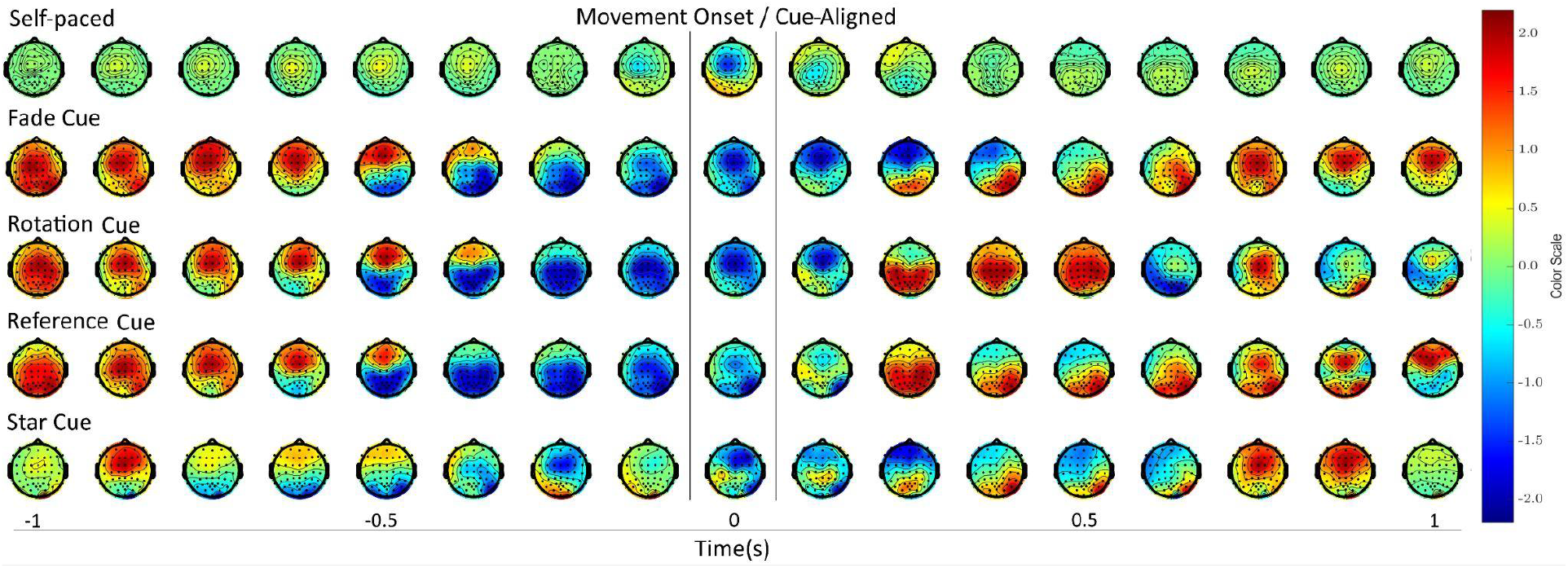
Topoplots comparison between self-paced (movement onset) and cue-based (aligned on cue onset) data. The time spans from -1 to 1 s, relative to the movement onset respectively cue-aligned. Each topographic plot is separated by a time span of 125 ms. All plots are standardized to a consistent color scale ranging from -2.2 to 2.2 μV.

**Figure 8:**
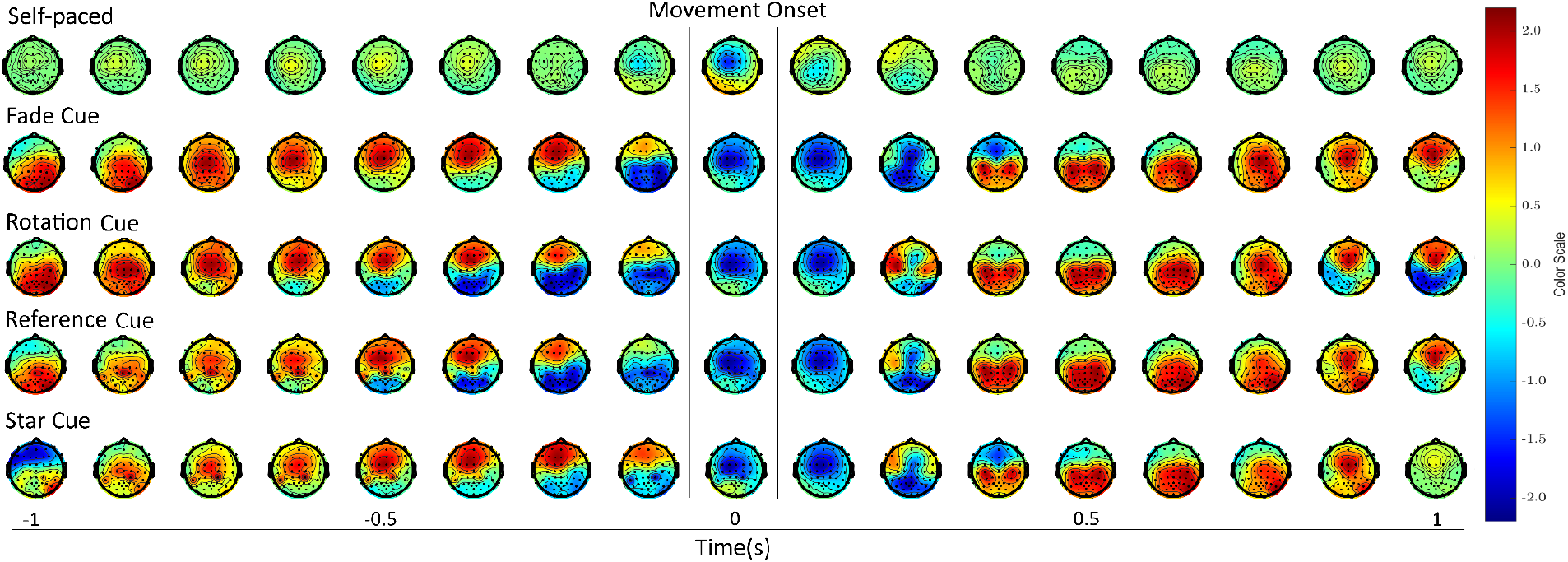
Topoplots comparison between self-paced and cue-based (triggered on movement onset) data. The time vector spans from -1 to 1 s, relative to the movement onset. Each topographic plot is separated by a time span of 125 ms. All plots are standardized to a consistent color scale ranging from -2.2 to 2.2 μV.

### Classification - movement vs rest

Fig. 9 depicts the time-varying cross-validated classification results for distinguishing between movement (encompassing all gestures) and no movement (rest), aggregated across all participants and based on different visual cues (cue aligned). Notably, the Rotation and Reference cue exhibited the highest accuracy, approximately 65.9% respectively 66.2%, at 0.97 s after the cues ‘go’ phase onset. This accuracy surpasses the real chance level (Mueller-Putz et al. 2008) of 54% for the two classes. The Fade Cue reaches 64.3% at 0.53 s and the Star Cue 63.6% at 0.33 s. In a parallel analysis on self-paced data triggered at the actual movement onset and comparing it to the same rest class, an accuracy of about 71.5% was achieved when averaged across all participants. When evaluating the classification results of the different cues with the self-paced data, as depicted in Fig. 10, it becomes evident that the Rotation and Reference cues exhibit only slightly inferior performance compared to the MRCPs generated from self-paced data. Conversely, the Fade and Star cue exhibit noticeably poorer performance. Additionally, the classification results differ when the MRCPs are aligned to the cue onset compared to the movement onset. However, for all cues, there is negligible variance between the movement onset for the different cue and the self-paced outcome.

**Figure 9:**
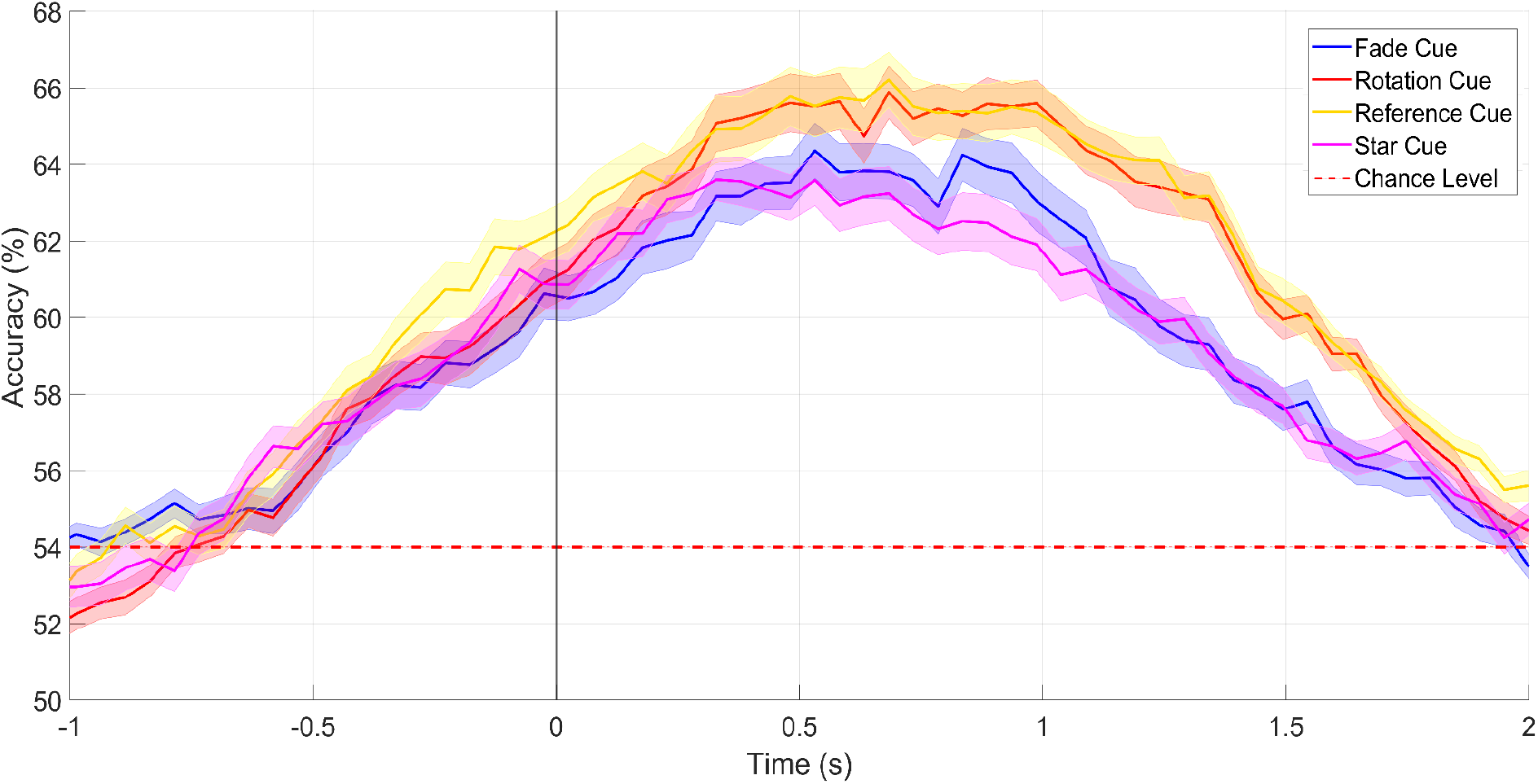
Classification results - cue aligned (movement vs rest) of the different cues (± standard error). Fade Cue (blue), Rotation Cue (red), Reference Cue (yellow), Star Cue (magenta), real chance level (red dashed).

**Figure 10:**
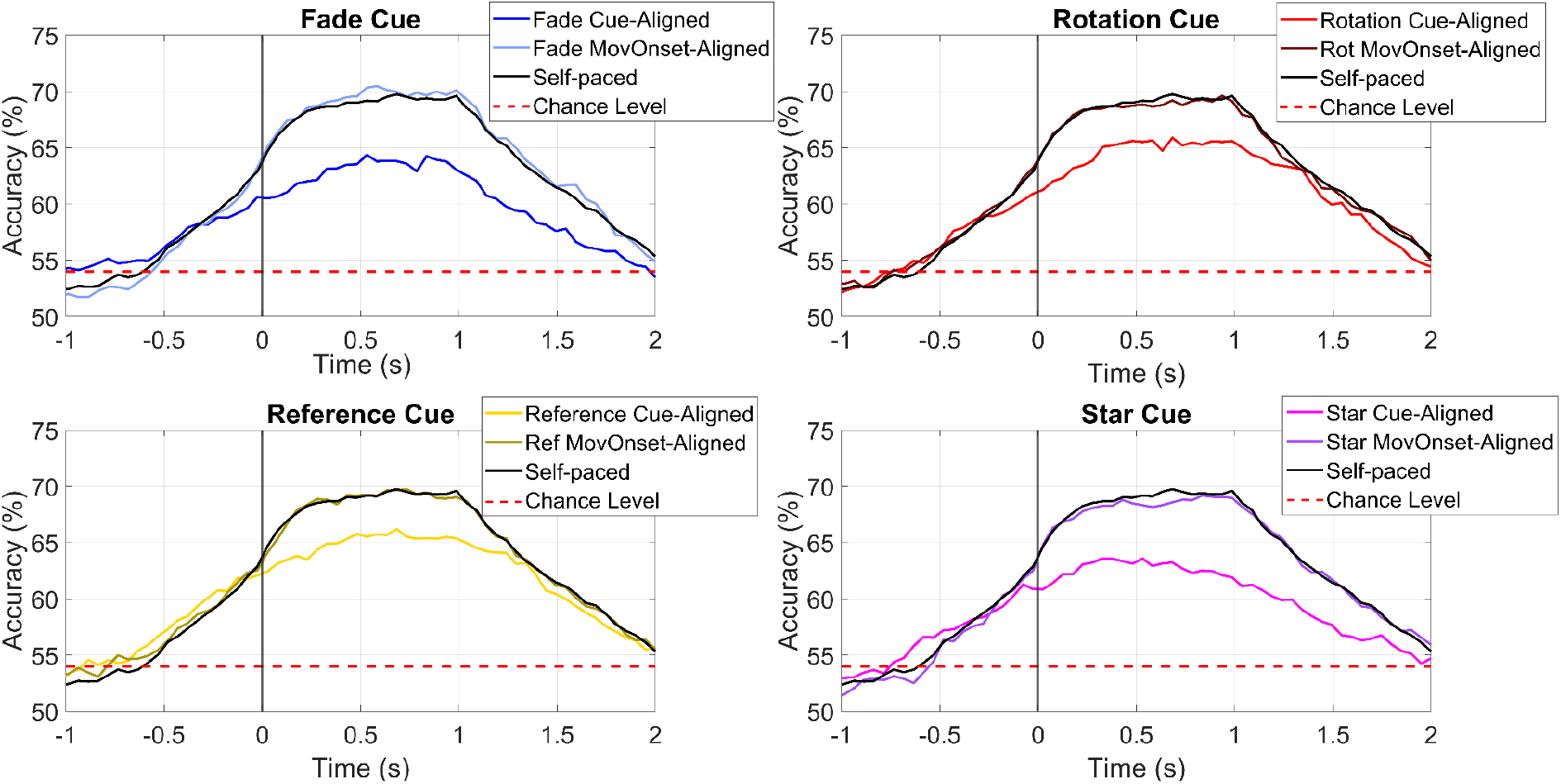
Classification results (movement vs rest) of the different cues (cue-aligned) compared to the movement onset and self-paced data. Fade Cue (blue), Rotation Cue (red), Reference Cue (yellow), Star Cue (magenta), real chance level (red dashed). The movement onset is plotted in a slight variation of the cue aligned color and the self-paced data is always plotted in black.

## Discussion

In this study, our aim was to reduce the impact of cues on MRCPs by introducing several types of visual cues for a cue-based data collection of movement. Classifiers derived from this data were applied on cue-less movement EEG. We have demonstrated that the data collection under the Rotation and Reference cue is minimally influenced by the cue effect, whereas data collected under the Fade and Star cue exhibit a noticeable higher degree of influence.

The timing variability results indicate that the Rotation and Reference cue provide a significantly more precise indication of when to initiate the movement compared to the Fade and Star cue. Consequently, we noted greater temporal variability in the onset of movements with the Fade and Star cue, resulting in blurred grand average potentials. These results underscore the significance of cue design, emphasizing the necessity for precise initiation timing of movements, while also minimizing VEPs that could affect the MRCPs during the transition from the cue’s start to the ‘go’ position. This is critically important in cases such as training patients with severe paralysis, where they need to refine their movement attempts for later use in a self-paced manner. (Ofner et al. 2019) introduced the concept of a gradually diminishing visual cue; however, no analysis was conducted to assess the precision of the cue in indicating movement.

The examination of grand average MRCPs in the time domain aligned to the different cue onsets reveals notable distinctions. Grand average MRCP shapes associated with the Reference and Rotation cue closely approximate those observed in self-paced data, whereas those for Fade and Star cue show minimal similarity. This observation aligns with findings regarding the timing variability of cues. Cues indicating movement onset more precisely replicate a self-paced approach when aligned to the cue onset. Considering the substantial similarity among the cues compared to self-paced data, it becomes evident that all cue types exhibit the most significant similarity until the start of the MP respectively after it. The non-existence of significant similarity during the MP of the MRCP can be attributed to the time shift of the cue onset between the cue-based data and the self-paced data’s movement onset. The objective is to minimize this shift to overlay the curves effectively and therefore increase the similarity for each timepoint. But for the training of a classifier this time shift can be easily addressed by adjusting the feature window. The spatial maps emphasize a conspicuous distinction between the self-paced data (triggered on movement onset) and the cue-based data (aligned on cue onset respectively movement onset). The Star cue data (aligned to cue onset) stands out prominently, exhibiting distinct characteristics compared to the other cue types which can possibly be explained by its smooth transition and its similar start position compared to the end position of the cue. On the other hand the Star Cue shows the least prominent MP pattern in the contralateral sensorimotor area around time 0 s. The MRCP patterns generated by the other three cue types exhibit mostly similar behavior, except for the time point at 0 s, where notable differences arise. Specifically, the Reference Cue displays a less pronounced negative amplitude and a narrower distribution around the motor area compared to the other two cue types. This can be explained by the temporal variability and that at this time point the MP is already over and the movement-monitor potential starts. When analyzing the cue-based data triggered solely on movement onset, a consistent pattern emerges across all spatial map time points, except for the -1 s and 1 s time points, which notably stand out for the Star Cue. This could also be explained by the unique design of the Star Cue and its smooth transition respectively similar start and end position which does elect less potential. The overall spatial pattern observed across all cues notably differs from the self-paced data. When training a classifier, it is crucial to consider this distinction and prioritize the selection of features from the cue-based data that are most similar to those observed in the self-paced approach.

The classification results demonstrate that movements signaled by the Rotation and Reference cue are more discriminable compared to the rest class, in contrast to the Fade and Star cue. This alignment reinforces the assumption that a precise cue indication of the starting time is important to closely resemble MRCPs, resulting in enhanced classification accuracy. Specifically, when comparing classification results to self-paced data triggered on movement onset, the accuracy values for the Rotation and Reference cue much closer mirror those of self-paced data, while Fade and Star cue exhibit lower accuracy. This observation underscores the effectiveness of the Rotation and Reference cue in closely approximating the accuracy achieved in self-paced scenarios, while highlighting the limitations associated with the Fade and Star cue.

## Conclusion

In this study, our findings demonstrate that the visual cue effect on MRCPs can be reduced to a minimum by introducing a new cue design. The timing variability analysis shows a more precise indication of movement onset with the Rotation and Reference cue. MRCP shapes associated with the Reference and Rotation cue resemble self-paced data much closer, while those triggered by the Fade and Star cue show much less similarity. Consistent with those findings, classification results reinforce the notion that movements signaled by Rotation and Reference cue are more discriminable compared to rest, in contrast to the Fade and Star cue. This suggests that a cue which evolves gradually over time and has a precise indication of the starting time leads to enhanced classification accuracy against rest. Specifically, when comparing classification results to self-paced data triggered on movement onset, the accuracy values for the Rotation and Reference cue closely mirror those of self-paced data, while the Fade and Star cue exhibit significantly lower accuracy. In the context of training a classifier, the highlighted characteristics of the Rotation and Reference cue emerge as pivotal factors contributing to the classifier’s efficacy, particularly in online scenarios. These findings underscore the significance of leveraging the precision embedded in visual cues, specifically Rotation and Reference cue, to attain accurate classifications. In a broader sense, the outcomes of this study emphasize the importance of gradual visual changes and the critical role of visual cue precision in ensuring dependable and distinguishable outcomes related to MRCPs.

## Acknowledgements

This project is funded by the European Union’s HORIZON-EIC-2021-PATHFINDER CHALLENGES program under grant agreement No 101070939 and by the Swiss State Secretariat for Education, Research and Innovation (SERI) under contract number 22.00198.

The authors especially acknowledge productive and beneficial discussions with the colleagues Markus Crell, Johanna Egger and Kyriaki Kostoglou.

## Notes

### Competing Interest Statement

The authors have declared no competing interest.

